# Topology-based Sparsification of Graph Annotations

**DOI:** 10.1101/2020.11.17.386649

**Authors:** Daniel Danciu, Mikhail Karasikov, Harun Mustafa, André Kahles, Gunnar Rätsch

**Affiliations:** Biomedical Informatics Group, Department of Computer Science, ETH Zurich, Zurich, Switzerland; Biomedical Informatics Research, University Hospital Zurich, Zurich, Switzerland; Swiss Institute of Bioinformatics, Zurich, Switzerland; Department of Biology, ETH Zurich, Zurich, Switzerland

## Abstract

Since the amount of published biological sequencing data is growing exponentially, efficient methods for storing and indexing this data are more needed than ever to truly benefit from this invaluable resource for biomedical research. Labeled de Bruijn graphs are a frequently-used approach for representing large sets of sequencing data. While significant progress has been made to succinctly represent the graph itself, efficient methods for storing labels on such graphs are still rapidly evolving. In this paper, we present RowDiff, a new technique for compacting graph labels by leveraging expected similarities in annotations of vertices adjacent in the graph. RowDiff can be constructed in linear time relative to the number of vertices and labels in the graph, and in space proportional to the graph size. In addition, construction can be efficiently parallelized and distributed, making the technique applicable to graphs with trillions of nodes. RowDiff can be viewed as an intermediary sparsification step of the original annotation matrix and can thus naturally be combined with existing generic schemes for compressed binary matrices. Experiments on 10,000 RNA-seq datasets show that RowDiff combined with Multi-BRWT results in a 30% reduction in annotation footprint over Mantis-MST, the previously known most compact annotation representation. Experiments on the sparser Fungi subset of the RefSeq collection show that applying RowDiff sparsification reduces the size of individual annotation columns stored as compressed bit vectors by an average factor of 42. When combining RowDiff with a Multi-BRWT representation, the resulting annotation is 26 times smaller than Mantis-MST.

## 1 Introduction

The exponential increase in global sequencing capacity [1] and the resulting growth of public sequence repositories have created an urgent need for the development of compact representation schemes of biological sequences. Such schemes should not only maintain all relevant biological sequence variation but also provide fast access for sequence search and extraction. After initial attempts focused on the lossless compression of full sequences, e.g., using the Burrows-Wheeler transform [2], the field soon turned towards representing a proxy of the input sequences instead: the sets of all *k*-mers contained in them. For this, any recurrent occurrence of a substring of length *k* in the input is represented by a unique *k*-mer, forming a *k*-mer set. A query of a given sequence against the input text can then be replaced by exact *k*-mer matching against the set. Longer strings are queried as a succession of *k*-mers. Although it is a lossy representation of the input (as, e.g., repeats longer than *k* are collapsed), constructing *k*-mer sets has proved highly useful in practice [3, 4, 5, 6].

### 1.1 Representation of *k*-mer sets

Various representations have been developed to balance the trade-off between the space taken by the *k*-mer set and query time or representation accuracy. Conceptually, the *k*-mer set fully defines a vertex-centric *de Bruijn* graph, where each *k*-mer forms a vertex and arcs are represented implicitly, based on whether any two vertices share a *k*−1 overlap. The simplest representations are bitmaps or (perfect) hash-tables that indicate the presence or absence of any possible *k*-mer over the input alphabet in the input text. While non-optimal in space, they offer constant-time query of *k*-mers. More compact representations use approximate membership query data structures to probabilistically represent a de Bruijn graph [7, 8] or utilize succinct de Bruijn graphs (a generalization of the Burrows-Wheeler transform) [9], which usually require less than one byte per input *k*-mer over the nucleotide alphabet {*A, C, G, T* }.

### 1.2 De Bruijn graph annotation

A major limitation of the above representations is that the identities of any sequence labels contained in the input text set are lost. To alleviate this, the concept of *colored de Bruijn graphs* emerged [10] (otherwise known as annotated or labeled de Bruijn graphs), allowing for the representation of additional annotations per *k*-mer. These annotations can either be stored in conjunction with the *k*-mers or be organized in a separate data structure, using the *k*-mer representation only as an index space. Although the first option is used by a number of conceptually interesting methods, such as Mantis [11] that uses counting quotient filters to represent the *k*-mers linked to an annotation identifier, here we will only focus on the second option, as it allows for the connection of arbitrary annotations to the *k*-mer set, without re-processing the *k*-mer index.

Conceptually, the set of annotations is a relation between *k*-mers and labels that can be represented as a binary matrix, where the *k*-mer set indexes the rows and each annotation label specifies a column. Any entry (*i, j*) in the matrix represents the relation of *k*-mer *i* and annotation *j*. Different methods have been suggested to compress this annotation matrix in a way that still allows for efficient query. VARI [12, 13] concatenates the rows of the annotation matrix and compresses the result using either an RRR [14] or Elias-Fano coding [15, 16].

Rainbowfish [17] takes advantage of high redundancy in matrix rows by computing a frequency code for the unique rows, compressing the unique rows in a matrix ordered by these codes, then representing the original matrix as a variable-length code vector. However, this method and other frequency coding-based approaches become less effective for data sets with greater levels of noise or inter-sample variability. Multi-BRWT [18] compresses the matrix in a hierarchical tree structure exploiting column similarity, but leaving the possible row redundancy unexploited. Alongside these methods, there is a rich literature of different compressors for graph annotations developed over the years, each improving on the compression performance of previous methods [19, 20, 21]. All of these methods share the common property that they act as general purpose binary matrix compressors, and thus, they do not take into account any particular domain knowledge in their construction.

### 1.3 Leveraging graph topology to improve annotation compression

While the methods mentioned above rely solely on similarities between annotation matrix elements to achieve their compression, a few have additionally leveraged graph topology to increase their compression potential. The Bloom filter correction method introduced by [22] encodes the columns of the annotation matrix in Bloom filters with high false positive rate. Assuming that all vertices within a graph unitig (a path in which all vertices except for the first and last have in- and out-degree 1) share identical annotations, a row in the annotation matrix (corresponding to all vertices from the same unitig in the graph) is computed as the bit-wise AND of the rows stored for every vertex of that unitig. While achieving high accuracy in decoding row annotations, the corrected Bloom filters are not able to losslessly decode the rows of the encoded annotation matrix. In addition, the authors introduce a lossless approach based on wavelet tries which leverages graph backbone paths to improve compression performance. However, these paths must be provided by the user and cannot be computed automatically by the method.

The more recently introduced Mantis MST method [23] constructs an annotation graph with nodes representing the unique rows of the annotation matrix. In this annotation graph, a weighted edge between two nodes *v*_1_ and *v*_2_ is created if there exist adjacent vertices *s*_1_ and *s*_2_ in the underlying de Bruijn graph whose annotations are represented by *v*_1_ and *v*_2_, respectively. The weight of this edge (*v*_1_, *v*_2_) is then set to the Hamming distance of the unique rows *v*_1_ and *v*_2_. Mantis MST computes the minimal spanning tree of the annotation graph and represents the annotation of a node as its bit-wise XOR with the annotation of its parent node in the spanning tree, while only the annotation of the root node is represented explicitly.

### 1.4 Our contribution

We present a new scheme for representing graph annotations, RowDiff, which takes advantage of similarities between the annotations of neighboring vertices to compress annotation matrices. RowDiff can be constructed using |*G*| + 3*m* + *O*(|***c***|) bits of memory, where |***c***| is the compressed size of the largest column in the annotation matrix and *G* is the size of the memory representation of the graph and in our case is less than 4*m* + *o*(*m*) bits [9], where *m* is the number of k-mers, thus making it suitable for annotating virtually arbitrarily large graphs. Since RowDiff is a transformation of the input annotation matrix attempting to increase its sparsity, RowDiff can be naturally chained with any generic scheme for compressed binary matrix representation to achieve further improvements in compression performance. We demonstrate the compression performance of RowDiff relative to the state-of-the-art lossless Rainbow-MST and MultiBRWT methods on datasets representing different annotation matrix densities.

In the next sections, we define the underlying concepts (Sections 2.1 and 2.2) and detail our methods for construction (Sections 2.3 to 2.4) and querying (Section 2.5) of the RowDiff data structures. We then describe the test datasets (Section 3.1) and study the representation sizes (Sections 3.2, 3.3, and 3.4), construction time (Section 3.5) and query time (Section 3.6) of RowDiff-compressed annotations. Finally, we discuss limitations and directions for future work (Section 4).

## 2 Method

### 2.1 Notation

We will operate in the following setting. Let *k* be a positive integer. The order *k* de Bruijn graph over a set of sequences *S*, denoted by DBG_*k*_(*S*), is a directed graph DBG_*k*_(*S*) := (*V*_*k*_, *E*_*k*_), whose vertices *V*_*k*_ are the set of all distinct sub-strings of length *k* of sequences in *S* (*k*-mers), and an arc links *u* ∈ *V*_*k*_ to *v* ∈ *V*_*k*_, if *u*_2:*k*_ = *v*_1:*k−*1_, where *s*_*i*:*j*_ denotes the sub-sequence of *s* from position *i* up to and including position *j*. We denote with deg^*−*^(*v*) and deg^+^(*v*), *v* ∈ *V*_*k*_ the in- and out-degree of a vertex, respectively. Vertices *v* ∈ *V*_*k*_, deg^*−*^(*v*) = 0 are called *source vertices* and vertices *v*∈ *V*_*k*_, deg^+^(*v*) = 0 are called *sink vertices*.

Given an arbitrary set of labels *L*, an *annotation* for a de Bruijn graph DBG_*k*_(*S*) is a relation 𝒜 ⊂ *V*_*k*_× *L*, which assigns to each vertex *v∈V*_*k*_ a set of labels, *l*(*v*) ⊂ *L*. We will trivially represent 𝒜 using a binary matrix 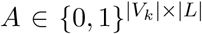, denote with *A*_*i*_ the *i*-th row of *A*, and with *A*_*i*_ ⊕*A*_*j*_ the element-wise XOR of rows *i* and *j*.

### 2.2 RowDiff transformation

RowDiff relies on the observation that adjacent vertices in the graph are likely similarly annotated, and thus, their respective rows in the annotation matrix *A* are similar as well. This implies that if (*u, v*) ∈*E*_*k*_, storing the difference between *A*_*u*_ and *A*_*v*_ may be more space efficient than storing *A*_*u*_, i.e. popcount(*A*_*u*_ ⊕ *A*_*v*_) *<* popcount(*A*_*u*_), where popcount(*x*) represents the number of set bits in row *x*.

RowDiff is defined as a transformation that converts an annotation matrix *A* of a de Bruijn graph into a new, sparser, annotation matrix *A*^*\**^ of the same size and an additional anchor vector 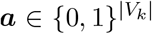, that is, 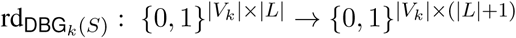. The anchor vector ***a*** stores which rows remain unchanged. We show that the original annotation matrix *A* can be reconstructed from the RowDiff transformed matrix *A*^*\**^ and the anchor vector ***a***. Empirically, the RowDiff transformed matrix is significantly better compressible in the typical case where neighboring vertices have similar annotations. We develop an efficient algorithm for defining good anchors and for computing the RowDiff transform 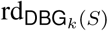 and its inverse.

For each vertex *u*∈ *V*_*k*_ we arbitrarily define its RowDiff successor as its lexicographically largest outgoing vertex succ(*u*), such that (*u*, succ(*u*)) ∈ *E*_*k*_ and succ(*u*) ≥ *v*∀ (*u, v*) ∈ *E*_*k*_, if such *u* exists. RowDiff replaces each row *A*_*u*_ with the (likely sparser) delta relative to its RowDiff successor. For binary rows, the delta is simply the element-wise XOR, 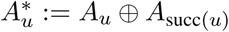,while for non-binary rows, the delta could store the difference between the row and its successor. In this work, we focus on binary matrices. The previous equation implies that 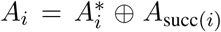,which gives us a simple formula for recursively reconstructing the original row. In order to be able to reconstruct the original annotation *A* from *A*^*\**^, some rows are left unchanged. A vertex *v* ∈*V*_*k*_ for which the annotation is stored unchanged is called an *anchor* and its corresponding value in the anchor bit vector will be set to 1, ***a***_*v*_ = 1. Sink vertices do not have a RowDiff successor, and must thus be anchors.

Algorithm 1 shows the implementation of the *inverse* transformation 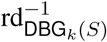, which reconstructs the original row *A*_*i*_ from the RowDiff representation *A*^*\**^.

**Algorithm 1** Row annotation reconstruction

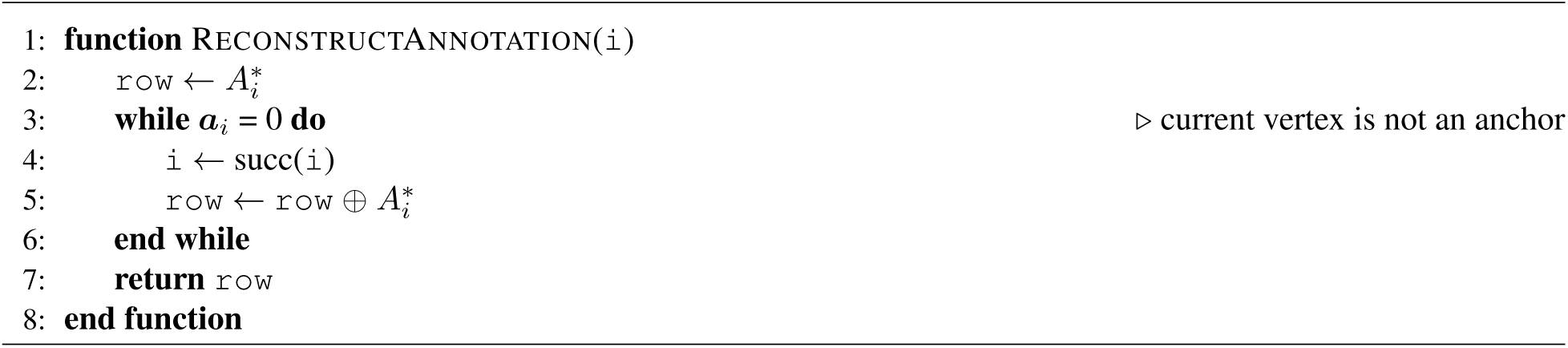

Starting from any vertex in the de Bruijn graph, Algorithm 1 defines a traversal leading to an anchor vertex, for which the annotation was not transformed. Since de Bruijn graphs may have cycles, additional anchor vertices might have to be assigned in order to break RowDiff cycles (those cycles where *every* vertex is a RowDiff successor relative to its predecessor in the cycle).

#### Proposition 1.

*Algorithm 1 finishes for every starting vertex, if and only if every sink vertex in the graph is an anchor and every RowDiff cycle contains at least one anchor vertex*.

*Proof*. Assume the algorithm does not finish for a starting vertex *i*. This implies that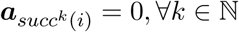, ∀*k* ∈ N. Since the number of vertices in the graph is finite, there must exist *l, m* ∈ ℕ, *l* ≠ *m*, s.t. *succ*^*l*^(*i*) = *succ*^*m*^(*i*). Thus, (succ^*l*^(*i*), succ^*l*+1^(*i*), …, succ^*m*^(*i*)) is a cycle and, hence, must contain at least one anchor vertex, which contradicts the initial assumption. Proof of necessity is equally trivial.

#### Proposition 2.

*Algorithm 1 correctly reconstructs the original annotation row A*_*i*_ *for every vertex i* ∈ *V*_*k*_. *Proof*. The algorithm computes 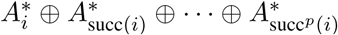, and thus, 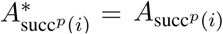. By repeatedly reducing the last 2 terms using 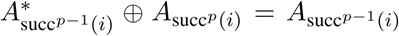,the original equation is reduced to *A*_*i*_, which is the desired value. □

Once the set of anchor vertices ***a*** satisfies Proposition 1, the RowDiff-transformed matrix *A*^*\**^ together with the anchor indicator bitmap ***a*** encode the original annotation matrix.

### 2.3 Anchor assignment

In addition to the small set of anchors described in Proposition 1, we seek to cap the maximum RowDiff path length (i.e. a path taken by Algorithm 1) to a certain value *M* (typically between 10 and 100) by ensuring that at least every *M* -th vertex in a RowDiff path is an anchor, as described below. This guarantees that the number of iterations in Algorithm 1 is bounded by a constant, and thus the average time complexity of reconstructing a single row is 𝒪 (*l. M*), where *l* ≪ |*L*| is the average number of set bits (labels) per row. At the same time, since anchor vertices require storing the original, less sparse annotation row, it is desirable to minimize the total number of anchor vertices in order to keep the popcount (and thus the compressed size) of the RowDiff annotation *A*^*\**^ small.

The following anchor assignment algorithm allocates anchor vertices near-optimally in four steps as follows (see Algorithm 2). First, we traverse RowDiff paths backwards (in parallel) starting from *sink vertices* (see Algorithm S1). The backward traversal stops either when we reach a source vertex or when we reach a vertex *v*∈ *V*_*k*_, s.t. succ(*v*) ≠ *u* for the previously traversed vertex *u* (see **Figure 1**, top). Note, the traversal is not terminated when reaching a vertex with multiple incoming arcs, but explores each of them and continues to further traverse these RowDiff paths backwards. When the distance from the current vertex to the next assigned anchor in the current RowDiff path reaches *M*, the vertex is marked as an anchor. In practice, once the backward traversal is finished, the vast majority of the vertices have been traversed, and the anchor assignment is optimal, in the sense that no anchors are closer than *M* to each other. In the second step, we start at *source vertices* and traverse RowDiff paths forwards, i.e. paths of the form *v*, succ(*v*), succ^2^(*v*), … (see Algorithm S2). The traversal stops when we reach an already visited vertex. In the third step, we start traversing forward at all forks with unvisited vertices. After the third step, the only vertices that were not traversed must belong to a simple cycle (a cycle where all vertices have deg^*−*^(*v*) = deg^+^(*v*) = 1). The fourth step traverses these cycles (in parallel). Each of these traversals sets an anchor every *M* vertices during the traversal. Since we visit each vertex only once, the time complexity of anchor assignment is 𝒪(|*V*_*k*_|).

**Figure 1:**
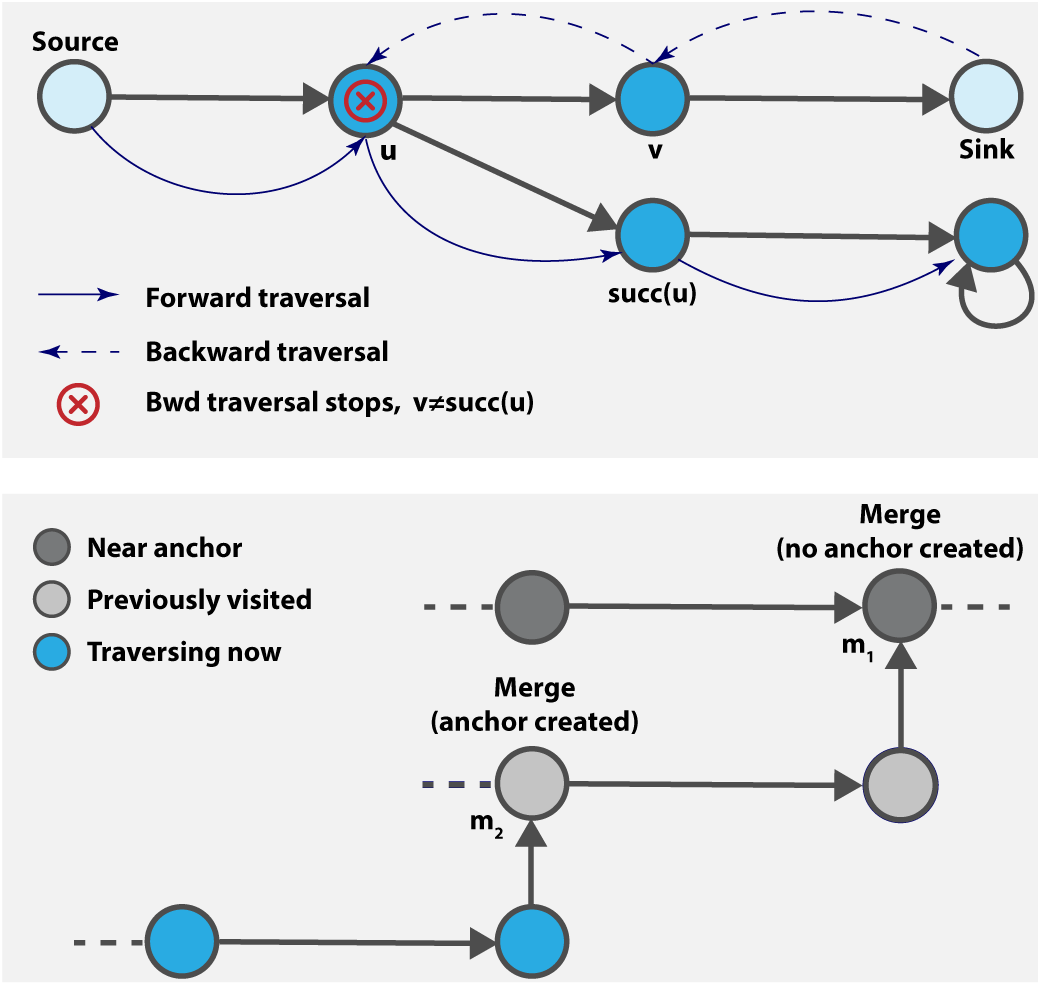
**Top:** RowDiff traversal. When traversing backward to assign anchor vertices, the traversal stops at vertex *u*, because succ(*u*)≠ *v*. When traversing forward, the last outgoing vertex is selected. **Bottom:** Chained merge. Dark grey vertices are marked as `nearAnchor`. When traversing the light grey vertices, we merge into *m*_1_, marked as `nearAnchor`, thus, no anchor is set. When traversing the blue vertices, an anchor must be set at *m*_2_, as *m*_2_ is not marked as `nearAnchor`.

**Algorithm 2** Anchor assignment

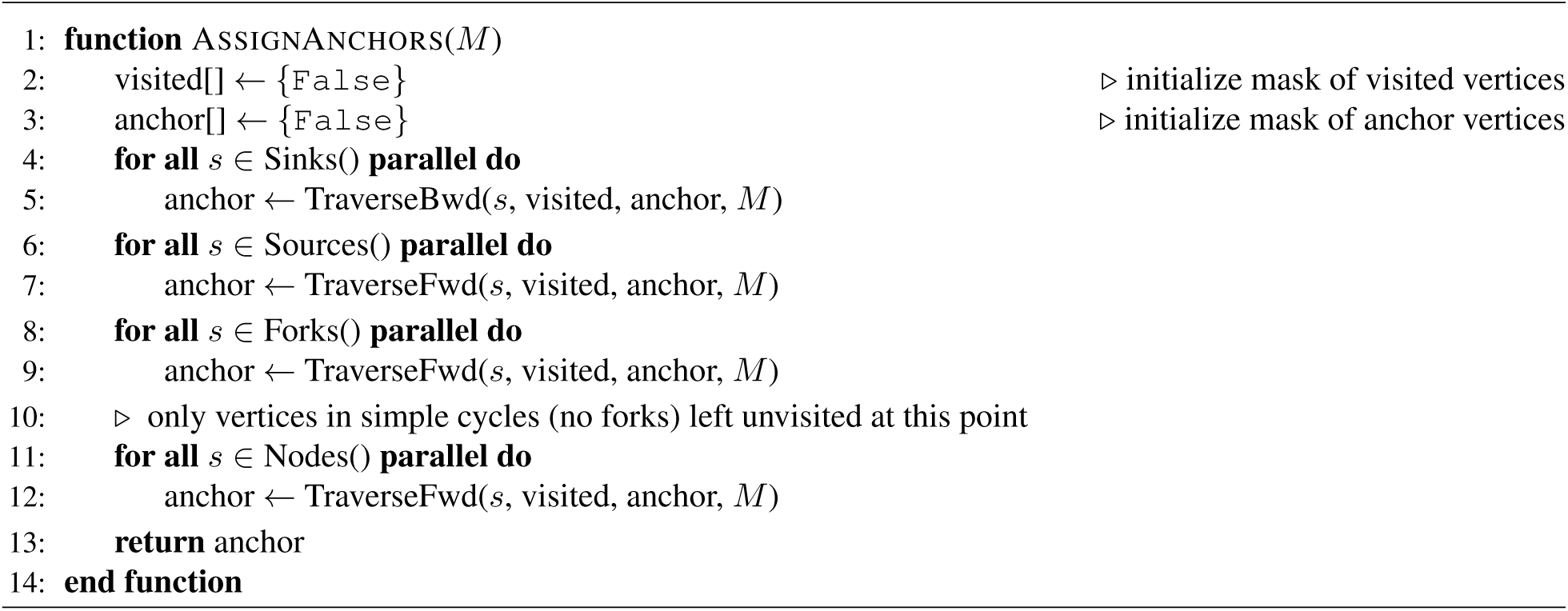

#### Proposition 3.

*The anchors assigned by Algorithm 2 guarantee successful termination of Algorithm 1 for any input vertex v* ∈ *V*_*k*_.

*Proof*. Step 1 of the algorithm trivially guarantees that all sink vertices are anchor vertices. Steps 3 and 4 guarantee that all cycles in the graph are traversed and at least one anchor vertex is set in each cycle. The conditions in Proposition 1 are thereby satisfied and Algorithm 1 finishes and successfully reconstructs *A* from *A*^*\**^.

One important detail in the forward traversal step is handling the situation when the traversal stops due to merging into a visited vertex. Not setting an anchor in such cases may result in arbitrarily long paths with no anchors (when such merges are chained). Always setting an anchor at a merge will introduce unnecessary anchors and increase the annotation density. We handle merges with the following simple heuristic: use an additional bit vector, `nearAnchor`, to mark all vertices that are known to be at a distance smaller than *M* to an anchor vertex. During forward traversal, when hitting a visited merge vertex not marked in `nearAnchor`, the anchor is not set (**Figure 1**, bottom). A more optimal algorithm for deciding if a merge vertex should create an anchor would require labeling each vertex with the distance to its nearest anchor. In our implementation we preferred the heuristic algorithm due to its significantly reduced space complexity.

#### 2.3.1 Anchor optimization

To guarantee that none of the rows 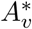 in *A*^*\**^ have more set bits than the corresponding row *A*_*v*_ in the original annotation, we perform the following anchor optimization procedure. For each *v* ∈ *V*_*k*_, s.t. popcount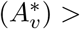 popcount(*A*_*v*_), we make such vertex an anchor, ***a***_*v*_ := 1, and replace 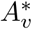 with *A*_*v*_. This ensures that all rows in the RowDiff-transformed annotation matrix are at least as sparse as the corresponding rows in the original annotation matrix.

##### Proposition 4.

*Each row in a RowDiff-transformed annotation matrix has the same or fewer set bits than its corresponding row in the original annotation matrix*.

The anchor optimization procedure is implemented similarly to the initial construction of RowDiff (see Section 2.4). Thus, it has the same time and space complexity.

### 2.4 RowDiff construction

A näive implementation of the RowDiff construction would be to load the matrix *A* in memory, and gradually replace its rows with their sparsified counterpart, while traversing the graph. Although fast and simple, this method requires to keep the entire annotation matrix *A* and the graph in memory. Unfortunately, often this is not realistic and even the annotation matrix *A* alone can easily reach several terabytes in size. Thus, we developed a distributed parallel construction algorithm that only loads a few columns of *A* at a time and, hence, needs a limited amount of memory.

In the first stage, we load the graph and for each vertex pre-compute the indices of the unique RowDiff successor and the (possibly multiple) RowDiff predecessors, stored in vectors `pred` and `succ`, respectively. The `pred` and `succ` vectors are used to build *A*^*\**^ in the second stage without the need to query the graph itself and load it in memory. To make the algorithm scale to de Bruijn graphs with trillions of vertices, vectors `pred` and `succ` are built and traversed in a streaming manner. They are loaded in small blocks, as described in Algorithms 3 and S3, and never kept in memory in full. Thus, pre-computing the `pred` and `succ` vectors essentially makes it possible to query the graph topology during the second stage while only using 𝒪(1) additional space. After the RowDiff annotation *A*^*\**^ has been generated, the `pred` and `succ` vectors are not required for querying and, thus, can be discarded.

**Algorithm 3** RowDiff transform

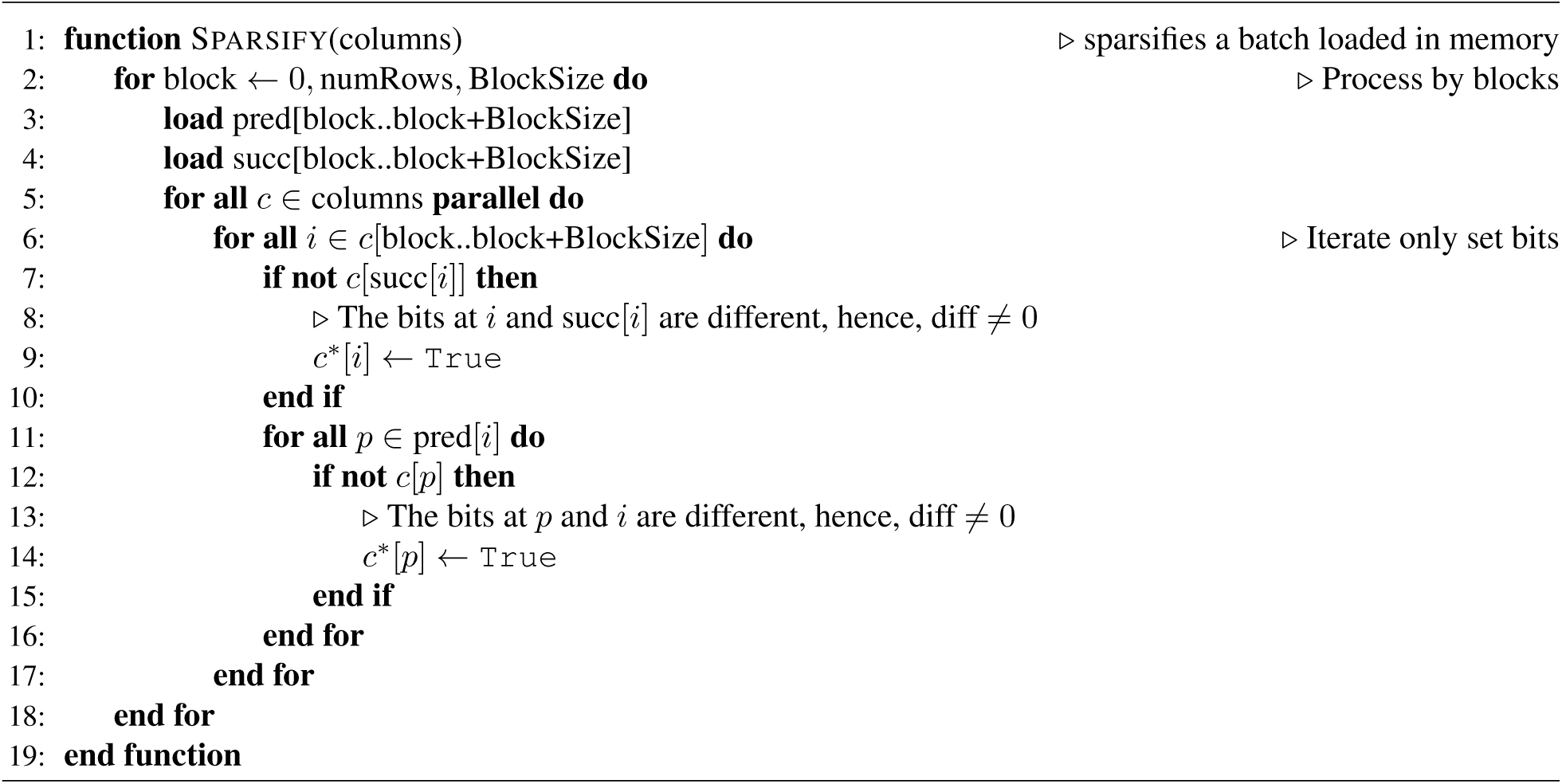

The second stage of the construction algorithm (the sparsification workflow) is schematically described in **Figure 2**. The initial sparsification of *A* can be trivially distributed by dividing the columns of *A* into groups and processing each group on a different machine. Each machine processes its assigned columns in batches. The size of each batch is determined dynamically by loading columns into memory until a desired upper limit is reached. This upper limit must be greater than the largest column being processed in compressed bit-vector format, but otherwise not restricted. For each column in the batch, we iterate only the set bits (only those rows corresponding to vertices annotated with the label represented by that column) and compare them with the bits at positions `pred` and `succ` in the same column to compute the RowDiff-transformed row, as shown in **Figure 2**.

**Figure 2:**
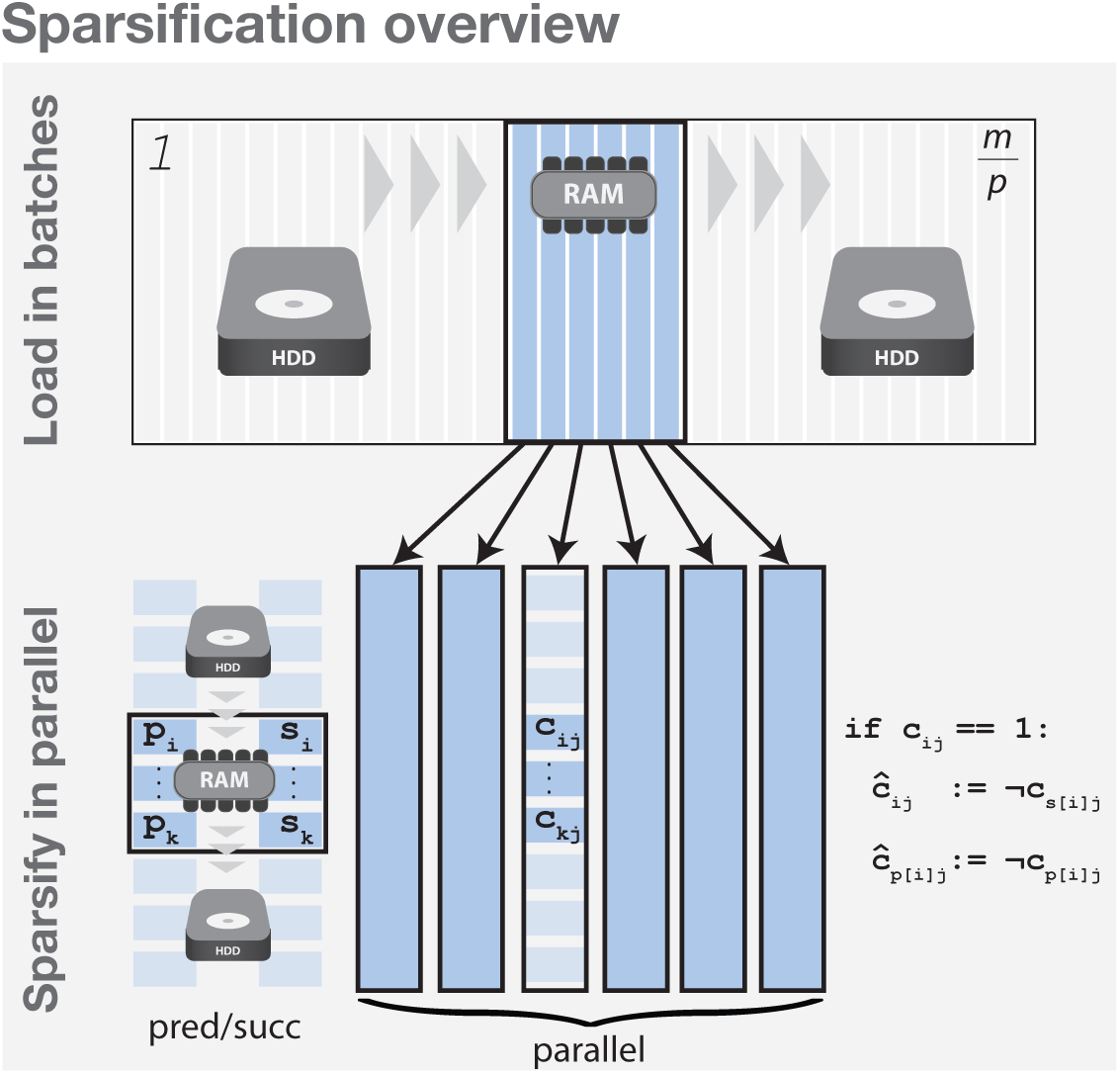
RowDiff transform algorithm – Schematic overview of sparsification on a single machine. **Top:** Columns are loaded into memory in batches (until memory is exhausted) and each batch is fully transformed to RowDiff. The result is serialized and the process moves on to the next batch. **Bottom:** Each batch is transformed to RowDiff as follows. The algorithm iteratively loads into memory blocks of the pre-computed vectors `pred` and `succ`. Then, all columns of the batch are processed in parallel. The algorithm iterates only through set bits of each column in the active block and computes the elements of the RowDiff transformed matrix *A*^*\**^ (see Algorithm 3 for a more detailed description).

#### 2.4.1 Scalability and complexity

Algorithm 3 only traverses set bits in *A*, and for each set bit in row *i* it performs 𝒪(deg^*−*^(*i*) + 1) operations, hence the total time complexity is 𝒪((1 + *α*)popcount(*A*)), where *α* is the average in-degree of the graph.

For de Bruijn graphs, *α* ≤ |Σ|, and hence the time complexity is linear in the number of set bits of the original annotation matrix, i.e. 𝒪 (popcount(*A*)). Algorithm S3, for constructing `pred` and `succ`, traverses each vertex exactly once, hence its time complexity is 𝒪 (|*V*_*k*_|). Since the buffer used by Algorithm S3 has a constant size, the space complexity is |DBG_*k*_(*S*) | + 𝒪(1), where |DBG_*k*_(*S*) | denotes the memory footprint of the graph, which, for instance, in the case of the BOSS representation [9], typically does not exceed 4*m* + *o*(*m*) bits, where *m* = |*V*_*k*_|. After taking into account Algorithm 2 for anchor assignment, which requires 3 additional bits per vertex to indicate anchors and the traversal state, and putting it all together, we get that the RowDiff-transform can be performed in (popcount(*A*) + |*V*_*k*_|) time and in |DBG_*k*_(*S*) | + 3| *V*_*k*_| + 𝒪 (|***c*** |) space, where ***c*** is the memory footprint of the largest (densest) column of *A* in a compressed bit-vector format. Note that the first term in the sum is usually the dominant.

In conclusion, we mention again that RowDiff construction can be easily distributed on multiple machines with modest hardware requirements and run in parallel on each machine, which makes the method very attractive for practical use on very large data sets.

### 2.5 Querying annotations for paths

We now note that, when querying annotations for paths in the graph, or sets of rows corresponding to vertices from a local neighborhood in the graph, Algorithm 1 leads to redundant reconstruction work, as many of the queried rows belong to the same RowDiff paths. To alleviate this, we perform the traversal first and pre-compute all RowDiff paths from the rows queried. Then, we query all diff rows in one batch and reconstruct annotations for each row from the query. This ensures that no arc in these paths is traversed more than once. Moreover, querying all rows in one batch often allows making the query of the underlying representation of the sparsified binary matrix faster by exploiting its potential intrinsic features (e.g., jointly querying *n* bits in *m* columns is more cache-efficient and faster than *n* queries of single bits in each of the *m* columns).

### 2.6 Implementation details

We implemented RowDiff as part of the MetaGraph framework [6]. The code for reproducing results of the experiments is available at https://github.com/ratschlab/row_diff. For storing original columns of the annotation matrix as well as the indicator bitmap with anchor vertices, we used the SD vectors from the sdsl-lite library [24] for compressed representation of bitmaps. For compression of the transformed annotation matrix, we used the Multi-BRWT representation scheme proposed in [18], with its improved and scaled up implementation from MetaGraph.

## 3 Results and Discussion

In this section, we evaluate the performance of the methods described above both in terms of their final representation sizes and their construction time. In addition, we also study the effect of the maximum RowDiff path length on the final RowDiff representation size of the compressed annotations. Finally, we evaluate the degree of size reduction that RowDiff provides on a per-column basis.

### 3.1 Data sets

We evaluated the compression performance of RowDiff on three data sets with different levels of sequence variability and thus graph density. Our first data set consists of all Fungi sequences from RefSeq release 97 [25], with annotations derived from the taxonomic IDs of the sequences’ respective organisms. Our second and third data sets are derived from the cohort of 10,000 publicly available human RNA-Seq experiments used in [23]. We constructed annotated de Bruijn graphs from the RNA-Seq data set in the same manner as in [23], using a *k* value of 23, albeit with two samples discarded due to their withdrawal from the Sequence Read Archive. We will refer to this data set as RNA-Seq (k=23). The third data set is constructed using the graph cleaning approach implemented in MetaGraph [6], using a *k* value of 31. We will refer to this data set as RNA-Seq (k=31). For evaluating construction time and representation size, we shuffled the samples in each data set and generated subsets of increasing size.

We evaluated RowDiff against MST [23], employed in Mantis [11], which, to the best of our knowledge, is the most compact annotation representation method to date. Similarly to Rainbowfish [17], MST reduces the original annotation matrix to a set of unique rows and consists of two components: a vector, mapping indexes of rows of the annotation matrix to its unique rows (color classes) and unique rows compressed in a minimum spanning tree. In Mantis, this mapping vector is included into a hash table storing the k-mers of the de Bruijn graph, which is usually at least an order of magnitude larger than the compressed annotation. Thus, to make a fair comparison, we exclude the large contribution of Mantis’ graph representation, and only consider the mapping vector, using the same representation as in Rainbowfish [17]. Thus, we refer to the MST annotation representation as Rainbow-MST. Note that Rainbow-MST forms a graph annotation representation which, similarly to RowDiff, can be used with any de Bruijn graph representation with indexed k-mers.

### 3.2 Representation size

We now compare the representation size for RowDiff and other state-of-the-art graph annotation compression methods.

Figure 3 shows the representation size for the RNA-Seq (k=31) and RefSeq (Fungi) data sets. On the RNA-Seq (k=31) data set, RowDiff-MultiBRWT effectively takes advantage of the topology of the graph annotation and the similarity of rows of the annotation matrix and achieves a nearly 4-fold size reduction compared to Multi-BRWT applied on non-sparsified columns. Compared to the Rainbow-MST method, RowDiff-MultiBRWT achieves a 2-fold size reduction. Rainbow-MST computation on the subsets with more that 4000 samples could not be computed because Mantis did not complete within the 10 day limit of our compute cluster. For this reason, we also plotted the size of the Rainbow-MST mapping vector, which, being a subset of the MST annotation data, represents a lower bound for Rainbow-MST.

**Figure 3.**
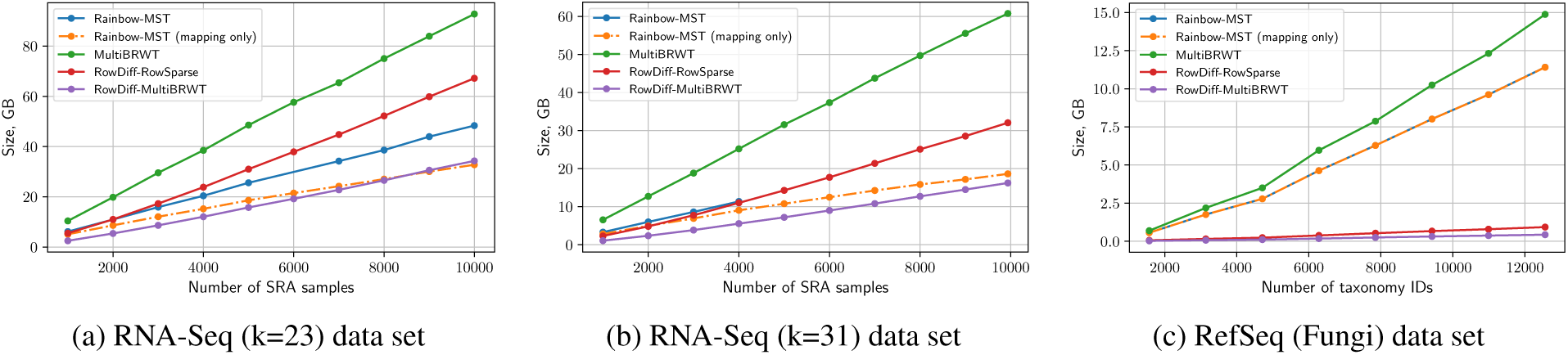
Representation size. The purple and red lines represent the size of the RowDiff annotation with and without MultiBRWT, respectively. The blue line indicates the size of the Rainbow-MST annotation. The orange line represents the size of the Rainbow-MST mapping vector and represents a lower bound on the Rainbow-MST representation size. Rainbow-MST computation on the RNA-Seq (k=31) data set with *>* 4000 samples did not complete within the 10 day limit of our compute cluster.

On the RefSeq (Fungi) data set, RowDiff takes advantage of the longer stretches of vertices with identical annotations and achieves a 26-fold size reduction relative to Rainbow-MST. Notably, this significant difference comes from the fact that virtually all of the space used by Rainbow-MST is taken by the mapping vector on this data set.

On the RNA-Seq (k=23) data set, RowDiff-MultiBRWT achieves a 2.5-fold size reduction relative to Multi-BRWT and a 1.5-fold reduction relative to Rainbow-MST.

The RowSparse format stores the indices of set bits in each row in a compressed integer vector. This type of annotation is faster to construct and to query than the Multi-BRWT representation, but its footprint is significantly larger on denser datasets, such as RNA-Seq (k=31).

### 3.3 Effects of graph density on compression

In this section we analyze how the density of the annotated graph affects RowDiff compression. In a first experiment we take a random subset of 1500 entries from the RefSeq Fungi data set and build graphs and corresponding annotations for k-mer sizes ranging from 15 to 31. **Table 1** shows how as the sparsity of the graph increases (with increasing k-mer length), the compression ratio |*A*|*/*|*A*^*\**^|sharply increases.

**Table 1:**
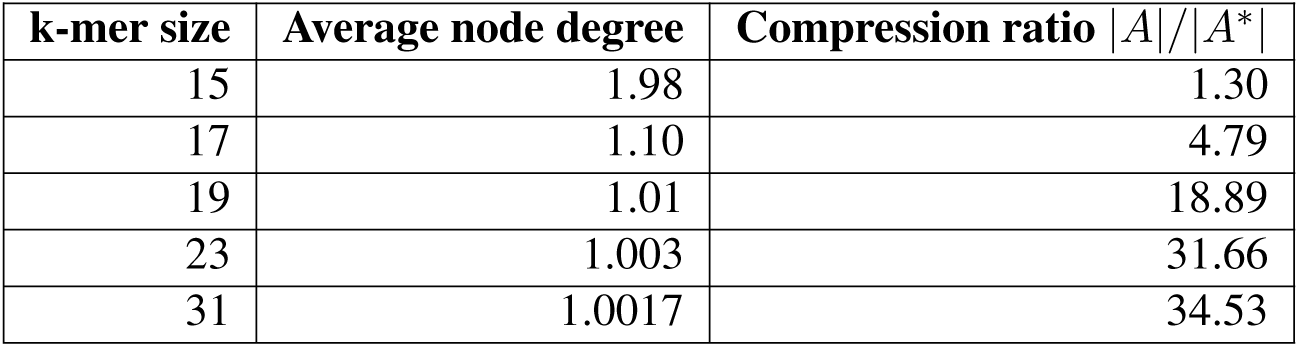
Compression ratio vs graph density. The sparser the graph the higher the compression ratio.

| k-mer size | Average node degree | Compression ratio $\vert A\vert /\vert A^*\vert $ |
| --- | --- | --- |
| 15 | 1.98 | 1.30 |
| 17 | 1.10 | 4.79 |
| 19 | 1.01 | 18.89 |
| 23 | 1.003 | 31.66 |
| 31 | 1.0017 | 34.53 |

In a second experiment, we test how the maximum path length *M* affects the annotation size for graphs of various densities. **Table 2** shows the annotation size on the RNA-Seq (k=23), RNA-Seq (k=31), and RefSeq (Fungi) data sets for various values of *M*. While increasing the maximum path length has negligible effect on the denser RNA-Seq graphs (with average node degrees of 1.08 and 1.04 respectively), it reduces the annotation size by a factor of up to 5.75 on the much sparser RefSeq (Fungi) graph (with an average node degree of 1.003).

**Table 2:** Annotation size vs maximum path length *M* for RNA-Seq (k=23,31) and Refseq Fungi (k=31). A sharp decrease in annotation size can be observed for the sparse RefSeq (Fungi) graph. Sizes shown in GB for RNA-Seq and MB for Refseq.

| Dataset | M=10 | M=20 | M=50 | M=75 | M=100 |
| --- | --- | --- | --- | --- | --- |
| RNA-Seq23 | 126 | 118 | 116 | 115 | 119 |
| RNA-Seq31 | 67 | 63 | 60 | 60 | 59 |
| Refseq Fungi | 1455 | 813 | 398 | 302 | 252 |

### 3.4 Compression of single columns

In this experiment, we measure how RowDiff compresses individual columns of the annotation matrix. **Figure 4** shows the compression factor 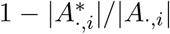 achieved by RowDiff on two datasets representing two different extreme cases of sequence variability.

**Figure 4:**
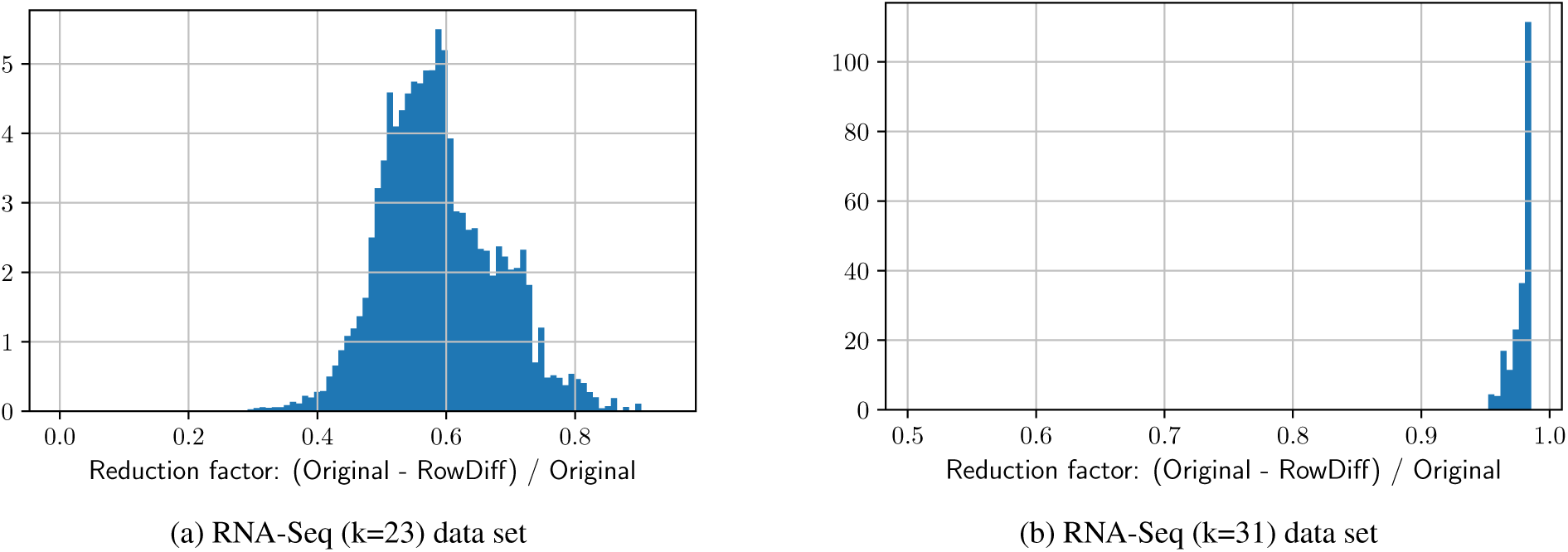
Histogram of the column reduction factor. 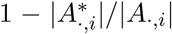.On the denser RNA-Seq (k=23) graph, the reduction factor peaks at 0.6, while on RNA-Seq (k=23) the reduction factor peaks at 0.97. The columns are stored as SD compressed vectors.

The de Bruijn graph constructed from assembled genomes RefSeq (Fungi) contains significantly fewer branches and bubbles than the graph constructed from reads RNA-Seq (k=31), thus its annotation is significantly better compressed by RowDiff, with an average reduction factor of 42.

### 3.5 Construction time

In **Figure 5**, we compare the construction times for building RowDiff and MST [23]. The construction time for RowDiff-MultiBRWT includes the RowDiff transform from original columns to the RowDiff format (with *M* = 100) in addition to the time for conversion of the transformed columns to the Multi-BRWT binary matrix representation. For MST, the time does not include construction of the mapping vector and includes only the time for compression of the unique annotation rows, which is a lower bound on the total construction time for the MST method. Note that the construction time for RowDiff-MultiBRWT grows linearly in the number of columns of the annotation matrix, and superlinearly for MST.

**Figure 5:**
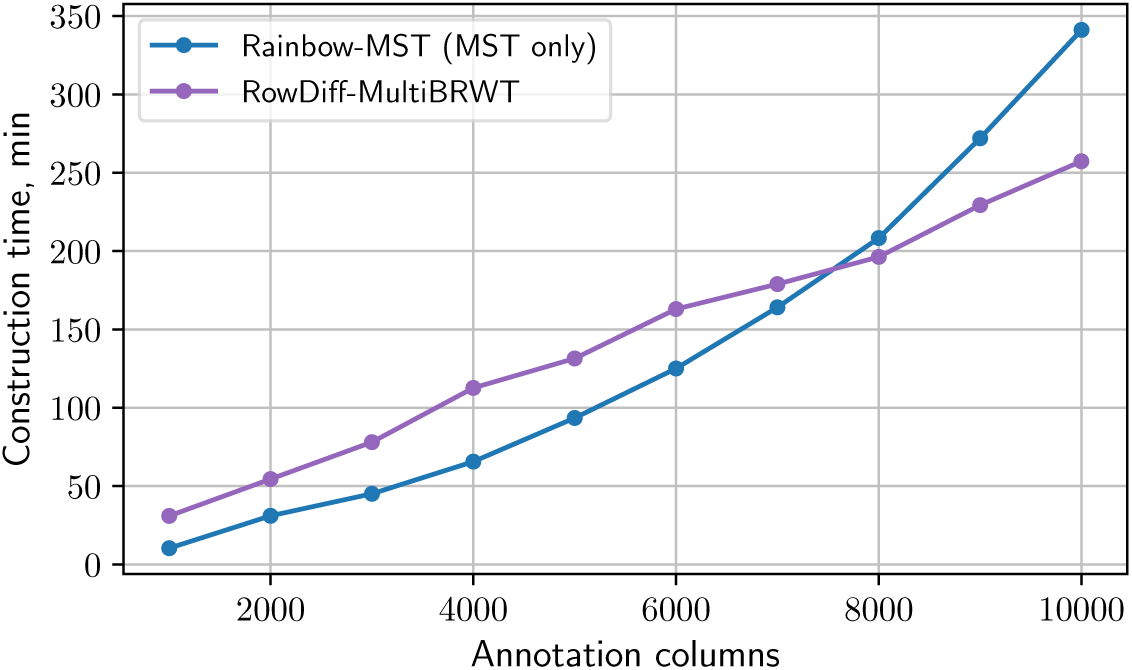
Construction time for the RowDiff and MST annotation representations on the RNA-Seq (k=23) data set with 72 threads.

### 3.6 Query performance

In this experiment, we measured the time needed for querying the RNA-Seq (k=23) annotation for human transcripts. The query is performed with the algorithm optimized for long paths (see Section 2.5). First, we construct a list of annotation rows that have to be reconstructed from the RowDiff format and a list of all diff rows for querying in the RowDiff matrix. Then, all these rows are queried and the original annotation rows are reconstructed. **Table 3** shows the time taken for reconstruction of the original annotations for 100 and 1000 random human transcripts, which includes the time for querying the diff rows and reconstruction of the original annotations. Since RowDiff requires traversing the de Bruijn graph to get RowDiff paths, the query time for RowDiff depends on the traversal performance of the underlying representation of the de Bruijn graph. In this experiment, we used the succinct de Bruijn graph representation available in MetaGraph [6].

**Table 3:**
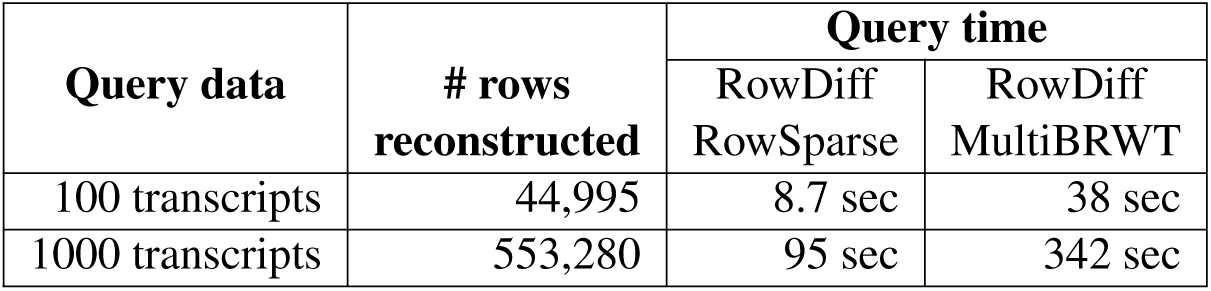
Time for querying 100 and 1000 random human transcripts with RowDiff-RowSparse and RowDiff-MultiBRWT. The second column shows the total number of original annotation rows reconstructed for the query. All benchmarks were performed with a single thread on Intel(R) Xeon(R) Gold 6140 CPU @ 2.30GHz.

## 4 Conclusions

In this paper, we introduced RowDiff, a new technique for compacting graph labels by leveraging the likely similarities in annotations of nodes adjacent in the graph. We designed a parallel construction algorithm with linear time complexity in the number of node-label pairs and small memory footprint. In addition, the algorithm can efficiently be distributed and parallelized, making it applicable on arbitrarily large graphs. RowDiff reduced the size of graph annotations by 2-to 26-fold when used in combination with Multi-BRWT relative to Mantis-MST, the most efficient state-of-the-art representation.

Although the row reconstruction method inevitably leads to an increase in *ad hoc* row query time due to the larger number of required annotation matrix queries, this limitation is alleviated in practice due to the tendency of real-world sequences to feature *k*-mers which co-occur on matching RowDiff paths.

The optimization of anchor assignment is a clear direction for future development of these methods. The anchor assignment method we have presented is designed to reduce the row reconstruction time by setting an upper bound on the traversal length. However, given that there is a trade-off between the size and the query time of the final representation, designing an objective function and a corresponding algorithm to best optimize these measures is a non-trivial task.

Moving beyond the representation of binary relations, a simple extension of the RowDiff method can be used as an efficient way to represent genomic coordinates for indexes of reference genomes. By representing a coordinate at each anchor node, the coordinates of all other nodes in that anchor’s corresponding RowDiff path can be computed via their traversal distance to the anchor.

Each improvement in the compression of sequence graphs and their associated annotations opens up further opportunities for their real-world applicability. When handling large annotations, even a 2-fold difference in the representation size can make a previously unapproachable annotation accessible to the available hardware. With RowDiff, we have demonstrated that there still is great potential for improving the representation of annotations on sequence graphs.

## Acknowledgements

Mikhail Karasikov and Harun Mustafa are funded by the Swiss National Science Foundation Grant No. 407540 167331 “Scalable Genome Graph Data Structures for Metagenomics and Genome Annotation” as part of Swiss National Research Programme (NRP) 75 “Big Data”. A. K. and D. D. are funded from ETH core funding to Gunnar Rätsch.

## Notes

### Competing Interest Statement

The authors have declared no competing interest.

## References

[1] Stephens, Z. D. et al. Big data: astronomical or genomical? PLoS biology 13, e1002195 (2015).

[2] Cox, A. J., Bauer, M. J., Jakobi, T. & Rosone, G. Large-scale compression of genomic sequence databases with the burrows–wheeler transform. Bioinformatics 28, 1415–1419 (2012).

[3] Ondov, B. D. et al. Mash: fast genome and metagenome distance estimation using minhash. Genome biology 17, 132 (2016).

[4] Breitwieser, F., Baker, D. & Salzberg, S. L. Krakenuniq: confident and fast metagenomics classification using unique k-mer counts. Genome biology 19, 198 (2018).

[5] Bradley, P., den Bakker, H. C., Rocha, E. P., McVean, G. & Iqbal, Z. Ultrafast search of all deposited bacterial and viral genomic data. Nature biotechnology 37, 152 (2019).

[6] Karasikov, M. et al. Metagraph: Indexing and analysing nucleotide archives at petabase-scale. BioRxiv (2020).

[7] Chikhi, R. & Rizk, G. Space-efficient and exact de bruijn graph representation based on a bloom filter. Algorithms for Molecular Biology 8, 22 (2013).

[8] Benoit, G. et al. Reference-free compression of high throughput sequencing data with a probabilistic de bruijn graph. BMC bioinformatics 16, 288 (2015).

[9] Bowe, A., Onodera, T., Sadakane, K. & Shibuya, T. Succinct de Bruijn graphs. In Lecture Notes in Computer Science (including subseries Lecture Notes in Artificial Intelligence and Lecture Notes in Bioinformatics) (2012).

[10] Iqbal, Z., Caccamo, M., Turner, I., Flicek, P. & McVean, G. De novo assembly and genotyping of variants using colored de bruijn graphs. Nature genetics 44, 226–232 (2012).

[11] Pandey, P. et al. Mantis: A Fast, Small, and Exact Large-Scale Sequence-Search Index. Cell Systems (2018). URL http://dx.doi.org/10.1016/j.cels.2018.05.021.

[12] Muggli, M. D. et al. Succinct colored de Bruijn graphs. Bioinformatics (2017).

[13] Muggli, M. D., Alipanahi, B. & Boucher, C. Building large updatable colored de bruijn graphs via merging. Bioinformatics 35, i51–i60 (2019).

[14] Raman, R., Raman, V. & Satti, S. R. Succinct indexable dictionaries with applications to encoding k-ary trees, prefix sums and multisets. ACM Transactions on Algorithms (TALG) 3, 43–es (2007).

[15] Elias, P. Efficient storage and retrieval by content and address of static files. Journal of the ACM (JACM) 21, 246–260 (1974).

[16] Fano, R. M. On the number of bits required to implement an associative memory (Massachusetts Institute of Technology, Project MAC, 1971).

[17] Almodaresi, F., Pandey, P. & Patro, R. Rainbowfish: A Succinct Colored de Bruijn Graph Representation. In Schwartz, R. & Reinert, K. (eds.) 17th International Workshop on Algorithms in Bioinformatics (WABI 2017), vol. 88 of Leibniz International Proceedings in Informatics (LIPIcs), 18:1–18:15 (Schloss Dagstuhl–Leibniz-Zentrum fuer Informatik, Dagstuhl, Germany, 2017). URL http://drops.dagstuhl.de/opus/volltexte/2017/7657.

[18] Karasikov, M. et al. Sparse binary relation representations for genome graph annotation. Journal of Computational Biology 27, 626–639 (2020).

[19] Bingmann, T., Bradley, P., Gauger, F. & Iqbal, Z. Cobs: a compact bit-sliced signature index. In International Symposium on String Processing and Information Retrieval, 285–303 (Springer, 2019).

[20] Harris, R. S. & Medvedev, P. Improved representation of sequence bloom trees. Bioinformatics 36, 721–727 (2020).

[21] Marchet, C. et al. Data structures based on k-mers for querying large collections of sequencing data sets. Genome Research 31, 1–12 (2021).

[22] Mustafa, H. et al. Dynamic compression schemes for graph coloring. Bioinformatics 35, 407–414 (2019).

[23] Almodaresi, F., Pandey, P., Ferdman, M., Johnson, R. & Patro, R. An efficient, scalable and exact representation of high-dimensional color information enabled via de bruijn graph search. In International Conference on Research in Computational Molecular Biology, 1–18 (Springer, 2019).

[24] Gog, S., Beller, T., Moffat, A. & Petri, M. From theory to practice: Plug and play with succinct data structures. In Lecture Notes in Computer Science (including subseries Lecture Notes in Artificial Intelligence and Lecture Notes in Bioinformatics) (2014).

[25] O’Leary, N. A. et al. Reference sequence (RefSeq) database at NCBI: Current status, taxonomic expansion, and functional annotation. Nucleic Acids Research (2016).

