## Supplemental Material for "Topology-based Sparsification of Graph Annotations"

### Supplementary Material
### Topology-based Sparsification of Graph Annotations

Daniel Danciu [1,2,*] Mikhail Karasikov [1,2,3,*] Harun Mustafa [1,2,3]
André Kahles [1,2,3,†] Gunnar Rätsch [1,2,3,4,†]

[1]Biomedical Informatics Group, Department of Computer Science, ETH Zurich, Zurich, Switzerland
[2]Biomedical Informatics Research, University Hospital Zurich, Zurich, Switzerland
[3]Swiss Institute of Bioinformatics, Zurich, Switzerland
[4]Department of Biology, ETH Zurich, Zurich, Switzerland

**Algorithm S1** Backward traversal for anchor assignment

```
function TraverseBwd(sink, visited[], anchors[], M)
    anchor[sink] ← True                              ▷ mark the sink as anchor
    queue.push(sink, 0)                              ▷ distance to next anchor is zero
    while not queue.empty() do
        node, depth ← queue.pop()
        if not visited[node] then                    ▷ for detecting loops
            visited[node] ← True
            if depth = M then
                anchor[node] ← True
                depth ← 0
            end if
            if Last(node) then                       ▷ only for last outgoing nodes
                for all n ∈ Incoming(node) do
                    queue.push(n, depth+1)           ▷ go further from anchor
                end for
            end if
        end if
    end while
end function
```

[*]Joint-first authors.

**Algorithm S2** Forward traversal for anchor assignment

```
function TraverseFwd(node, visited[], anchor[], nearAnchor[], M)
    path ← []
    while not visited[node] do                              ▷ Traverse until hitting a merge
        visited[node] ← True
        path.append(node)
        node ← lastOutgoing(node)
    end while
    for i = 0, len(path) − M, M do                          ▷ Assign anchor every M nodes
        anchor[path[i]] ← True
        nearAnchor[path[i:i+M]] ← True
    end for
    next ← lastOutgoing(path.back())
    if len(path) mod M == 0 or nearAnchor[next] then
        return                                              ▷ We merged close to an anchor, done
    end if
    anchor[path.back()] ← True
    nearAnchor[path[i:]] ← True
end function
```

**Algorithm S3** Streaming construction of the `pred` and `succ` vectors

```
function CreatePredSucc
    for start = 0, |V|, BLOCK_SIZE do            ▷ traverse the graph in blocks of size BLOCK_SIZE
        parfor i = start, start+BLOCK_SIZE do
            if a[i] = 0 then                                ▷ Anchors don't have a successor
                succBuf[threadNo].append(lastSucc(i))
            end if
            for all j in pred(i) do
                if succ(j)=i then                           ▷ Check if i is the RowDiff successor of j
                    predBuf[threadNo].append(j)
                    predBoundaryBuf.append(0)
                end if
            end for
            predBoundaryBuf.append(1)
        end parfor
                                                ▷ the following commands dump the memory buffer to disk
        succ.append(succBuf), pred.append(predBuff), predBoundary.append(predBoundaryBuf)
    end for
end function
```
